# *Tspecies*, Rapid Optimization for Estimating Species Divergence Time Using *K*_s_ Distribution

**DOI:** 10.1101/2025.05.18.654758

**Authors:** Mi-Jia Li, Xiao-Xue Li, Lin-Lin Xu, Bo-Wen Zhang

## Abstract

*K*_s_ distribution, the distribution of the synonymous substitutions, has been widely used to estimate the species divergence using orthologous genes. However, conventional approaches often ignore the underlying bias that species divergence is delayed to average gene divergence by 2*N*_e_ generations, where *N*_e_ represents the ancestral effective population size, due to the lack of scalable methods for *N*_e_ inference. Here, we demonstrate through simulations that *K*_s_ distribution variance correlates with *N*_e_, enabling direct estimation of ancestral population parameters from standard *K*_s_ data. Leveraging this relationship, we present *Tspecies*, an R package that corrects divergence time estimates using only substitution rates and *K*_s_ distributions, without requiring additional genomic data. Our practical application of *Tspecies* in *Liriodendron* has inferred a divergence time between North American and East Asian lineages (3.49 Ma) that align with late Pliocene cooling, and a large ancestral *N*_e_ (∼5 × 10^5^) consistent with fossil evidence. By incorporating a readily estimated *N*_e_, our tool resolves a long-standing bias in *K*_s_-based dating while maintaining computational efficiency and broad applicability.

*Tspecies* is freely available under an MIT license at https://github.com/limj0987/Tspecies.git.

## Introduction

*K*_s_ distribution, the distribution of the synonymous substitutions of orthologs, has been widely employed to estimate the species divergence times in comparative genomics. When studying the characteristics of some specific genes in *Gossypium hirsutum* (Hao *et al*. 2020), *Raphanus sativus* (Hu *et al*. 2018), *Brassica napus* (Zhu *et al*. 2020), researchers have successfully employed the *K*_s_ distributions to depict the divergence times of the objects from their close relatives. The *K*_s_ value reflects the sum of independent evolutionary distances accumulated in two species following their divergence. Building on this, under the assumption of constant mutation accumulation rates in both species, the divergence time *T* can be calculated by dividing the *K*_s_/2 value (representing the distance from either species to their common ancestor) by the substitution rate (*μ*). This *K*_s_ distribution-based method for estimating species divergence times has been widely adopted in comparative genomic studies and can be easily obtained with the commonly used genomic analysis toolkits such as OrthoFinder (Emms & Kelly. 2019) and KaKs_Calculator (Zhang. 2022).

However, under the coalescent theory framework, when post-divergence gene flow is not considered, the genetic divergence time between orthologs always predates the species divergence time (Sigwart. 2009). Therefore, accurate estimation of species divergence time requires calculating the difference *dT* between these two temporal scales. And for *dT*, according to the coalescent theory, the species divergence time differs from the average gene divergence time by 2*N*_e_ (Fig. 1A), where *N*_e_ represents the ancestral effective population size (Wakeley & Sargsyan. 2009). Current methods for inferring *N*_e_ primarily rely on population genomic data to reconstruct historical demographic dynamics (Charlesworth. 2009). However, due to limitations in sampling or sequencing, most studies lack sufficient population-level genomic data to estimate *N*_e_ (Wang *et al*. 2016). To address these constraints, alternative approaches that can estimate both species divergence and *N*_e_ using single or few genomes have thus been developed, such as the Bayesian Phylogenetics & Phylogeography (BPP) program (Yang. 2015) and the F1-hybrid Pairwise Sequentially Markovian Coalescent (hPSMC) model (Cahill *et al*. 2016). Nevertheless, methods like BPP face significant challenges: high computational demands, strict data requirements (e.g. high quality or haplotype-resolved genome), and reliance on assumptions of evolutionary parameters (e.g. random mating, independent recombination blocks). Their reliability further diminishes under challenges from polyploids (Yan *et al*. 2022) or ghost introgressions (Pang & Zhang. 2024). Therefore, this *N*_e_-dependent bias is often overlooked in comparative genomic studies because of the difficulty of obtaining ancestral *N*_e_. Directly scaling *N*_e_ through *K*_s_ distributions could therefore offer a more feasible solution for a general inference of species divergence times in comparative genomic studies. Of note, the practical application of our tool on the genomes from genus *Liriodendron* indicated a divergence time of ∼3.5 million years ago (Ma) between the North American and East Asian lineages, which gives a more plausible scenario of allopatric speciation driven by glacial divergence.

**Figure 1.**
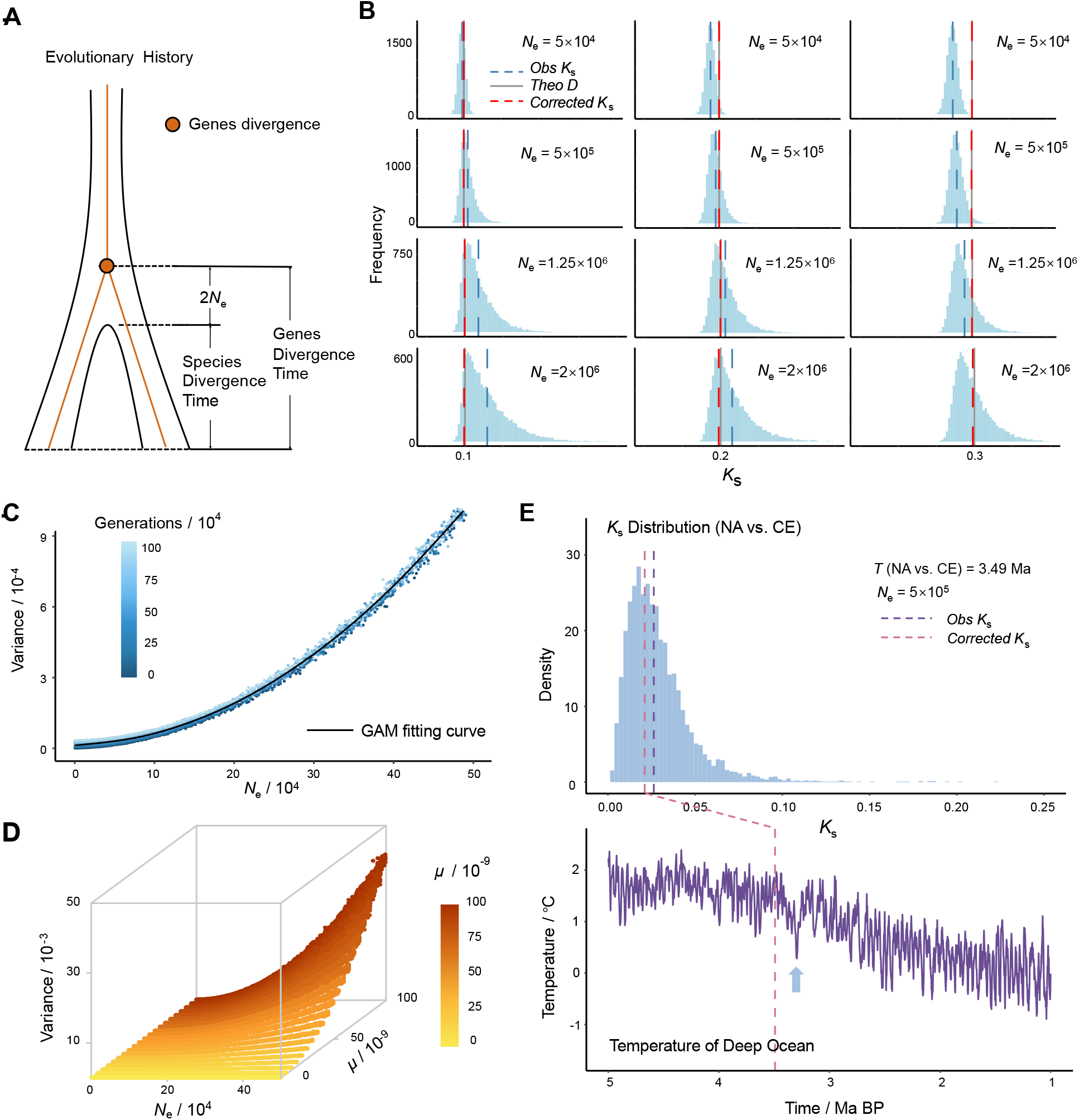
The optimization for the distribution of the synonymous substitutions (*K*_s_) **(A)** Schematic diagram of species divergence time and the average divergence time of genes. **(B)** Observed, theoretical, and corrected *K*_s_ in the *K*_s_ distribution of different simulated scenarios. The *Obs K*_s_ refers to the *K*_s_ directly calculated by the sequential differences; The *Theo D* refers to the theoretical species distance, which equals to twice *μ* multiplied by the simulated species divergence time; The *Corrected K*_s_ refers to the estimated species divergent distance corrected with 2*N*_e_ and multiple substitution. **(C)** The Generalized Additive Model (GAM) describing the relationship between *N*_e_ and variance of *K*_s_ distribution. **(D)** 3D Plot of Variance of *K*_s_ distribution vs. *N*_e_ and *μ*. **(E)** *K*_s_ distribution *(Top)* for orthologs between *L. tulipifera* from North America (NA) and *L. chinense* from eastern China (CE), and the paleotemperature trends *(Bottom)* over the past ∼5 Ma (before 2005) derived from the Lisiecki and Raymo (2005). Divergence time between NA and CE inferred by *Tspecies* is marked with pink dashed line; The sharp decline of global average temperature at ∼3.3 Ma is marked by a blue arrow.

## Results and discussion

The core concept underlying this optimization is to induce the coalescent theory, where in a population with a constant effective population size of *N*_e_, the expected time and the standard deviation for two alleles from a random locus to coalescence is 2*N*_e_ (Fig. 1A). Considering that the *K*_*s*_ distribution reflects the loci divergence from two sister species, these loci should only be coalescent in the ancestral population and thus the variance of the *K*_*s*_ distribution should also be positively correlated with ancestral *N*_e_. Therefore, we simulated the species divergence and modeled the *N*_e_-*K*_s_ variance relationship.

Simulation analyses revealed critical dependencies between *N*_e_ and *K*_s_-based divergence estimates. As illustrated in Fig. 1B, direct species divergence estimation using mean *K*_s_ values showed systematic deviations proportional to *N*_e_. However, applying *N*_e_-correction through multi-locus substitution models (Nielsen & Slatkin. 2013) reduced these deviations, aligning gene-level divergence with true species divergence times. Crucially, we established that the variance of *K*_s_ distributions serves as a quantitative proxy for *N*_e_, though this relationship exhibited phase-specific modulation during speciation events.

To dissect how species divergence time shapes *K*_s_ variance dynamics, we systematically varied *T* across 10^4^–10^6^ generations while maintaining a constant neutral mutation rate (*μ*). Fig. 1C illustrates the resultant *N*_e_-*K*_s_ variance relationships, where data points are color-coded by divergence time. Three salient patterns emerged: (1) a robust positive correlation between *N*_e_ and *K*_s_ variance persists across all *T* values under invariant *μ*; (2) at shorter divergence time,temporally distinct clusters (non-overlapping color groups) reveal *T*-dependent modulation of the relationship; (3) as *N*_e_ exceeds 2 × 10^4^, all trajectories converge toward the unified GAM curve (black), indicating diminishing temporal influence. This convergence highlights *N*_e_ dominance over *T* in governing variance saturation, consistent with coalescent theory predictions.

The generalized additive model (GAM) analysis revealed a strongly nonlinear yet highly predictive relationship between *N*_e_ and *K*_s_ variance (*R*^2^= 0.997, Fig. 1C). The smooth term for *N*_e_ showed significant complexity (effective degrees of freedom [edf] = 7.11). Model diagnostics confirmed robustness: the intercept (3.68, standard error = 0.0045) remained stable across divergence times (*t* = 815.8), while the nearly identical generalized cross-validation (GCV) score (0.0289) and scale estimate (0.0287) ruled out overfitting. Critically, this framework retains practical utility for estimating *N*_e_ from empirical *K*_s_ distributions, provided adequate sampling of gene orthologs across genomic regions.

Theoretical simulations demonstrate that *μ* significantly modulates the *N*_e_-*K*_s_ variance pattern (Fig. 1D). Higher *μ* values intensify the positive correlation between *N*_e_ and *K*_s_ variance, making *μ* an indispensable scaling factor for time calibration. Furthermore, as an established parameter in molecular clock dating, *μ* values are routinely estimated through fossil calibrations or cross-species comparisons, aligning with practical requirements for divergence time estimation. Therefore, we incorporated *μ* as a critical input parameter in our R package *Tspecies*.

While our model provides a computationally tractable framework for divergence time estimation, it adopts the simplifying simulation of identical sequence length and constant *N*_e_ across speciation events. To address the robustness of the observed *N*_*e*_-*K*_*s*_ Variance Relationship, we conducted sensitivity analyses under different sequential length (*L*) and post-divergence *N*_e_ variations. The relative difference in predicted variance in different *L* (Fig. S1) suggested potential sequence length effects at shorter *L* (e.g. *L* = 500bp), while longer lengths (*L* ≥ 1, 000bp) demonstrated more stable predictions (mean deviation = 12.2%). Additionally, assessments of subpopulation dynamics highlighted potential influences from post-divergence *N*_e_ variations (Fig. S2). However, the relative accuracy across different simulated scenarios indicated an acceptable bias of approximately 5.6% from the true divergence time, with higher *μ* yielding greater accuracy (Fig. S3).

Our practical application focusing on species from the genus *Liriodendron*, which are believed to have undergone significant population reduction throughout their evolutionary history (Chen *et al*. 2019). We compared *L. tulipifera* from North America (NA) with *L. chinense* from eastern China (CE) and from western China (CW) using their *K*_s_ distributions. The divergence time inferred by *Tspecies* is 3.49 Ma between NA and CE, and 3.53 Ma between NA and CW, which aligns the climate cooling (Lisiecki & Raymo. 2005) and ice sheet growth from the late Pliocene to the Ice Ages of the Pleistocene (Fig. 1E). Such abrupt climatic shifts are known to drive species range contractions and southward migration. Here, this cooling event may have fragmented ancestral *Liriodendron* populations by restricting suitable habitats, ultimately driving geographic isolation between North American and Asian lineages. In contrast, direct divergence estimates between *L. tulipifera* and *L. chinense* based on mean *K*_s_ values were older, reaching 4.36 Ma (NA vs. CE) and 4.40 Ma (NA vs. CW), tracing back to the warmer Mid-Pliocene period. Additionally, *Tspecies* estimated the *N*_e_ of the ancestral *Liriodendron* to be approximately 5 × 10^5^, which is consistent with the fossil evidence indicating that the genus was once widely distributed across the Northern Hemisphere (Ian Milne. 2006).

## Conclusion

By precomputing an ancestral *N*_e_ and scaling the gene divergence to species divergence under the coalescent theory framework, *Tspecies* fully utilized *K*_s_ distribution estimated from orthologous sequences. This package ensures optimal correction for species divergence time with dual consideration of biological realism and computational practicality.

## Materials and Methods

### Simulating the Divergence

To model the divergence of the two closely related species, we employed the *ms* (Hudson. 2002) and the *seq-gen* (Rambaut & Grass. 1997). Given that the samples were generated under neutral model, the rate of nucleotide substitutions rate between sequences was equivalent to the rate of synonymous substitutions. In each simulation, 10,000 pairs of homologous sequences of 1000 bp in length were generated, and the mean and variance of the *K*_s_ distribution were calculated using R.3.6.0. To account for the range of divergence scenarios, the substitution rate (*μ*) was set to 10^−7^ to 10^−10^, the theoretical divergence time (*T*) was set to 10^4^ to 10^6^ generations, and *N*_e_ was set to 500 to 500,000. These values are covered by θ and *t* in *ms*. In this case, *θ* is defined as 4*N*_e_*μ*, and *t* is the theoretical divergence time divided by four times the effective population size.

### Modeling the *N*_*e*_ Effects

Because the divergence time has little effect on the variance, we use *μ* as a categorical variable, and use Generalized Additive Models (GAMs) to analyze the *N*_e_-*K*_s_ variance relationship, selected for two reasons: **(a) Nonlinear Dynamics**, the relationship between two variables displayed strong nonlinearity, invalidating conventional linear models (e.g. LM). **(b) Unspecified Nonlinear Form**, without theoretical prior for the nonlinear pattern, prespecifying functional forms (as required by Nonlinear Least Squares, NLS) was impractical. GAMs circumvent this limitation through flexible smooth functions (e.g. splines) that adaptively capture nonlinear relationships. Based on the range of common *μ*, we divided different gradients and stored the corresponding data in R packages *Tspecies*, which were uploaded to GitHub for download.

### Robustness on *N*_e_-*K*_s_ Variance Relationship

To evaluate the impact of sequential length on the *N*_e_-*K*_s_ variance relationship, we compared model predictions across *L* = 500bp, 1,000bp, 1,500bp and 2,000bp while holding other parameters constant (*μ* = 10^−8^). For each *L*, we generated orthologous sequence pairs using *ms* and calculated the normalized variance of *K*_s_ distributions (Fig. S1).

In order to ascertain whether the predicted outcomes of the *Tspecies* undergo substantial alteration when *N*_e_ fluctuates, a simulation was conducted in *ms*: subsequent to species divergence, the number of one of the subpopulations diminished precipitously to one-half, one-fifth, and one-tenth of the ancestral population, respectively (Fig. S2). *Tspecies* was utilized to calculate the species divergence times, and the predicted outcomes were compared with the set theoretical times in simulation. Then, the relative accuracy (*RA*) was calculated by the following formula:

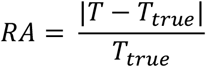

where *T* is species divergence time calculated by *Tspeices, T*_*true*_ is theoretical species divergence time. The relative accuracy under different *μ* are shown in Fig. S3.

### Application in *Liriodendron*

To test the model, we applied *Tspecies* to the genomes of genus *Liriodendron* with two distinct species from East Asian (*L. chinense*) and eastern North American (*L. tulipifera*). Genomes were downloaded from the database PRJNA418360 of NCBI. The *K*_s_ distribution of reciprocal best-hit gene pairs across the genomes was calculated using the *K*_s_ analysis pipeline implemented in the wgd package (Zwaenepoel *et al*. 2019). The obtained divergent distance is scaled with the reported synonymous substitution rate of 3.02 × 10^−9^ per site per year (Cui *et al*. 2006). The divergence time directly calculated by mean of the *K*_s_ distribution was compared to the time estimated under *Tspecies*. The *δ*^18^O data from the Lisiecki and Raymo (2005) are converted to direct temperature estimates using the equations of Hansen (Hansen *et al*. 2013).

## Authors’ contributions

M.L. and X.L. implemented the software package, performed simulation and validations, wrote the initial manuscript. L.X. helped with the practical application and validation. B.Z. contributed to study set-up, interpreted the results, and revised the manuscript. All authors read and approved the final manuscript.

## Competing interests

The authors have declared no competing interests.

## Acknowledgments

This project is supported by the National Undergraduate Innovation and Entrepreneurship Training Program and the Fundamental Research Funds for the Central Universities. Special thanks to Prof. Da-Yong Zhang for raising this pressing issue in current *K*_s_ estimation.

## Supplementary materials

**Figure S1.**
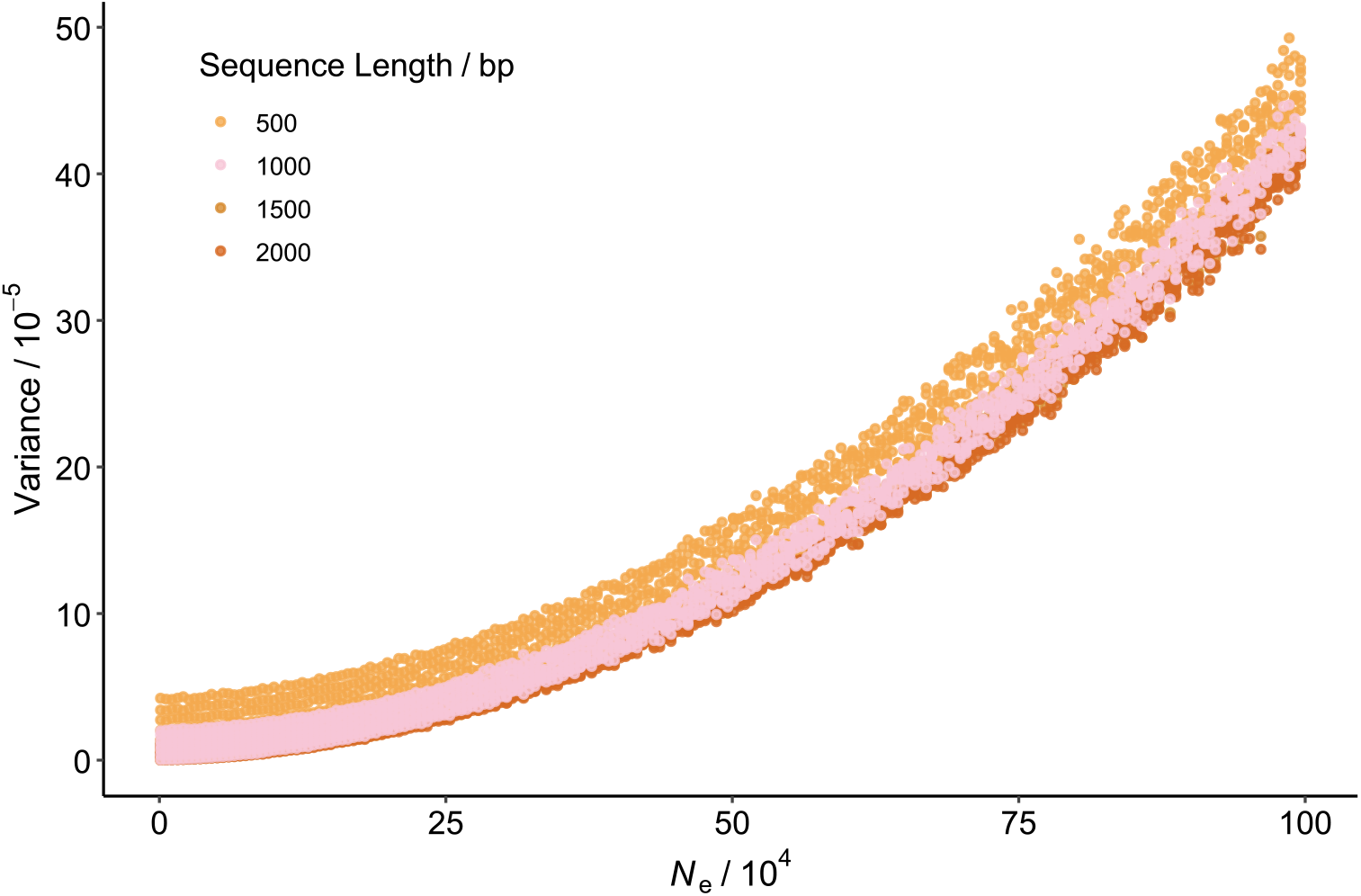
The relationship between *N*_e_ and variance of *K*_s_ distribution under different sequence length (*L*). The positive correlation patterns between *N*_e_ and *K*_s_ variance remained stable across tested *L* values (500, 1000, 1500, 2000bp). The relative difference suggested potential sequence length effects at shorter *L* (e.g. *L* = 500bp), while longer lengths (*L* ≥ 1, 000bp) demonstrated more stable predictions (mean deviation = 12.2%).

**Figure S2.**
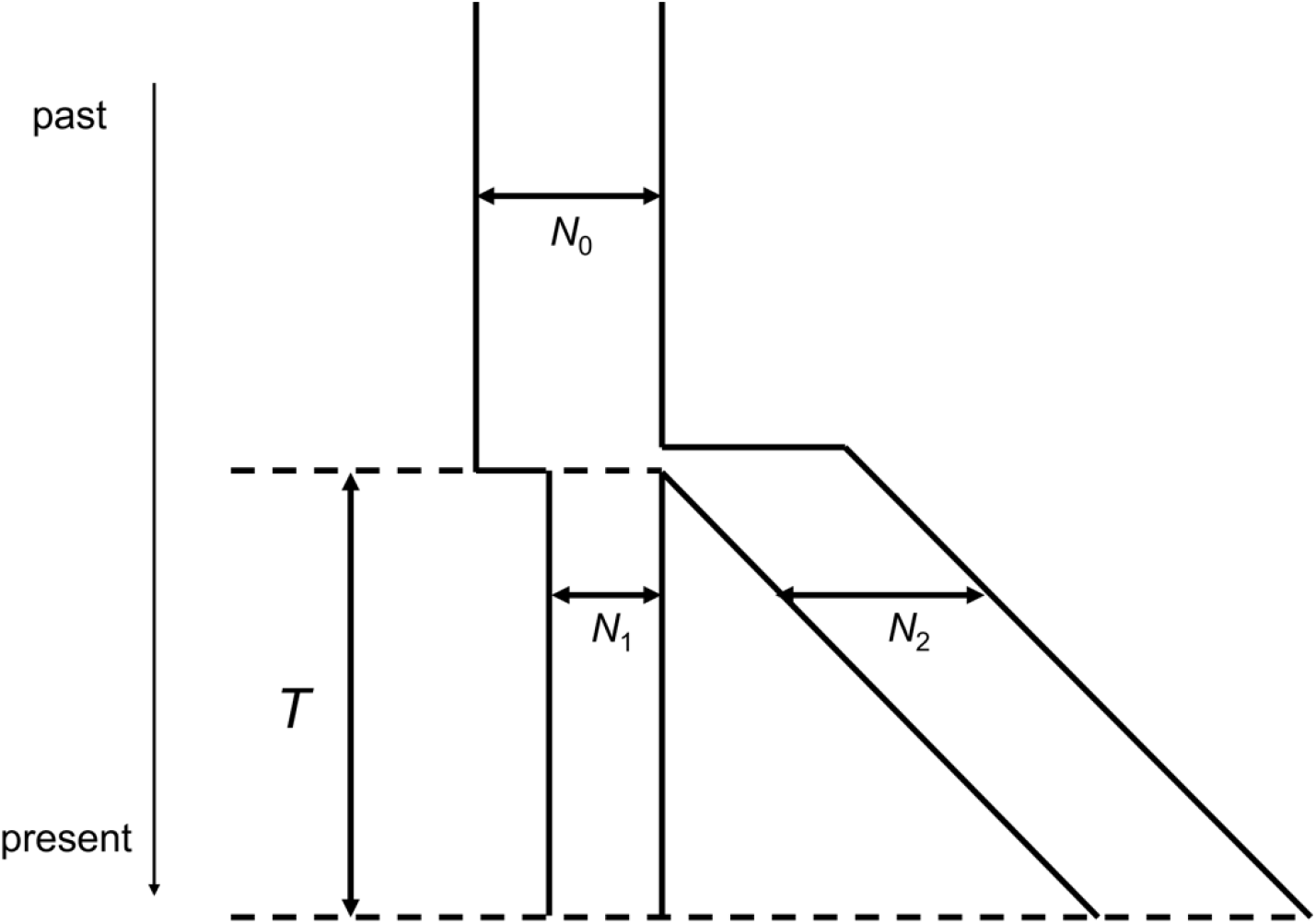
Schematic representation of species divergence scenarios. In simulation, the speciation event transpired *T* generations ago. Prior to the occurrence of speciation, the effective population size of the ancestor was *N*_0_. Immediately subsequent to the event of speciation, the effective population size of one of the subpopulations underwent a transition to *N*_1_, which corresponded to half, fifth, and tenth of *N*_0_, respectively. The effective population size of another remained constant and equal to that of the ancestor.

**Figure S3.**
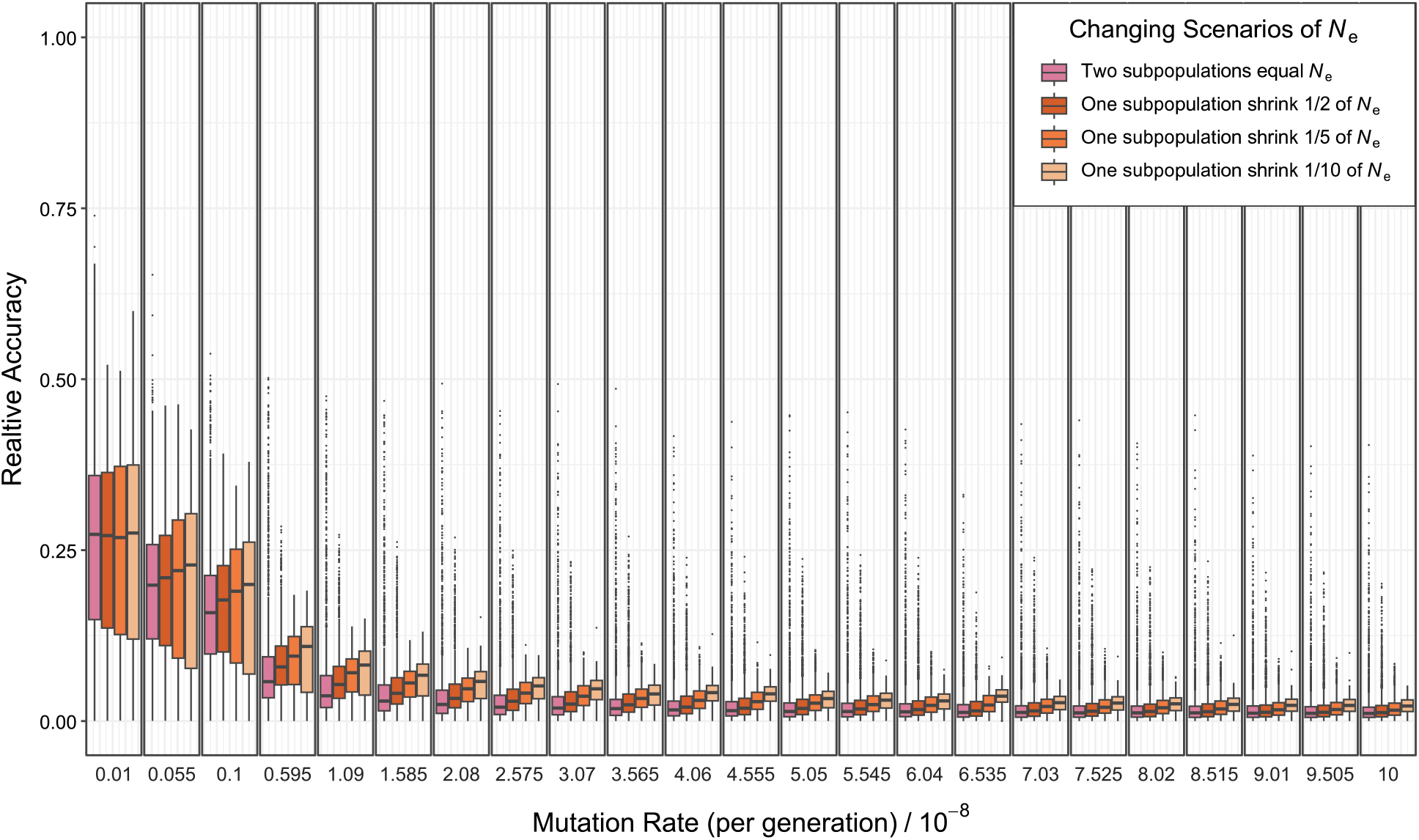
Relative accuracy of species divergence scenarios was calculated using simulated data. Yellow gradients represent scenarios where one subpopulation shrinks to 1/2, 1/5, or 1/10 of the *N*_e_, while the pink bar indicates no change in subpopulation size. We maintained constant subpopulation sizes in this study as the associated impact on accuracy was deemed acceptable (mean relative accuracy = 5.6%), with higher mutation rates (*μ*) showing improved scenario reconstruction accuracy. The *μ* gradient was consistent with Fig. 1D, as was the case for the final model (*Tspecies*).

## References

Cahill JA, Soares AER, Green RE, Shapiro B. Inferring species divergence times using pairwise sequential Markovian coalescent modelling and low-coverage genomic data. Philosophical Transactions of the Royal Society B: Biological Sciences 2016.

Charlesworth B. Effective population size and patterns of molecular evolution and variation. Nature Reviews Genetics 2009, 10(3) 195–205.

Chen J, Hao Z, Guang X, Zhao C, Wang P et al. Liriodendron genome sheds light on angiosperm phylogeny and species–pair differentiation. Nature Plants 2019, 5(1) 18–25.

Cui L, Wall PK, Leebens-Mack JH, Lindsay BG, Soltis DE et al. Widespread genome duplications throughout the history of flowering plants. Genome Research 2006, 16(6) 738–749.

Emms DM, Kelly S. OrthoFinder: Phylogenetic orthology inference for comparative genomics. Genome Biology 2019, 20(1) 238.

Hansen J, Sato M, Russell G et al. Climate sensitivity, sea level and atmospheric carbon dioxide. Philosophical Transactions of the Royal Society A: Mathematical, Physical and Engineering Sciences 2013, 371(2001) 20120294.

Hao P, Wang H, Ma L, Wu A, Chen P et al. Genome-wide identification and characterization of multiple C2 domains and transmembrane region proteins in Gossypium hirsutum. BMC Genomics 2020, 21(1) 445.

Hu T, Wei Q, Wang W, Hu H, Mao W et al. Genome-wide identification and characterization of CONSTANS-like gene family in radish (Raphanus sativus). PLOS ONE 2018, 13(9) e0204137.

Hudson RR. Generating samples under a Wright–Fisher neutral model of genetic variation. Bioinformatics 2002, 18(2) 337–338.

Ian Milne R. Northern hemisphere plant disjunctions: a window on tertiary land bridges and climate change. Annals of Botany 2006, 98(3) 465–472.

Lisiecki LE, Raymo ME. A Pliocene-Pleistocene stack of 57 globally distributed benthic d18O records. Paleoceanography 2005, 20(1).

Nielsen R, Slatkin M. An introduction to population genetics: Theory and applications. Sinauer Associates Sunderland, MA. 2013.

Pang XX, Zhang DY. Detection of ghost introgression requires exploiting topological and branch length information. Systematic Biology 2024, 73(1) 207–222.

Rambaut A, Grass NC. Seq-Gen: An application for the Monte Carlo simulation of DNA sequence evolution along phylogenetic trees. Bioinformatics 1997, 13(3) 235–238.

Sigwart J. Coalescent theory: An introduction. Oxford University Press. 2009.

Wakeley J, Sargsyan O. Extensions of the coalescent effective population size. Genetics 2009, 181(1) 341–345.

Wang J, Santiago E, Caballero A. Prediction and estimation of effective population size. Heredity 2016, 117(4) 193–206.

Yan Z, Cao Z, Liu Y, Ogilvie HA, Nakhleh L. Maximum parsimony inference of phylogenetic networks in the presence of polyploid complexes. Systematic Biology 2022, 71(3) 706–720.

Yang Z. The BPP program for species tree estimation and species delimitation. Current Zoology 2015, 61(5) 854–865.

Zhang Z. KaKs_Calculator 3.0: Calculating selective pressure on coding and non-coding sequences. Genomics, Proteomics & Bioinformatics 2022, 20(3) 536–540.

Zhu W, Wu D, Jiang L, Ye L. Genome-wide identification and characterization of SnRK family genes in Brassica napus. BMC Plant Biology 2020, 20(1) 287.

Zwaenepoel A, Van de Peer Y. Wgd-simple command line tools for the analysis of ancient whole-genome duplications. Bioinformatics 2019, 35(12) 2153–2155.

